# CD95/Fas Apoptosis Signal Initiation Depends on the Ligand Oligomerization State and Formation of Small Ligand-Receptor Complexes

**DOI:** 10.64898/2026.05.07.723507

**Authors:** Xiaoyue Shang, Nina Bartels, Nicolaas van der Voort, Andreas Neusch, Noah Salama, Gajen Thaventhiran, Ralf Kü hnemuth, Suren Felekyan, Claus A. M. Seidel, Cornelia Monzel

**Affiliations:** 2nd Institute of Physics, University of Stuttgart, Pfaffenwaldring 57, 70569 Stuttgart, Germany; Molecular Physical Chemistry, Heinrich-Heine University Düsseldorf, 40225 Düsseldorf, Germany; Experimental Medical Physics, Heinrich-Heine University Düsseldorf, 40225 Düsseldorf, Germany

**Keywords:** CD95, qSTED nanoscopy, apoptosis, IZ-CD95L, Fas, FasL

## Abstract

The death receptor cluster of differentiation 95 (CD95 / Fas) is an important inducer of apoptotic activity in tumor cells, but may exhibit differential cell responses (ranging from predominantly apoptosis up to proliferation) when stimulated by its CD95 ligand (CD95L/FasL). How ligand form and receptor–ligand stoichiometry shape these divergent outcomes remain unresolved. To define structural and functional determinants of CD95 signaling, we systematically compared the oligomerization and activity of native CD95L with CD95L variants, where ligand trimers were stabilized either by a FLAG-tag for subsequent monoclonal antibody (mAb) crosslinking or by genetically fused isoleucine zippers (IZ). Apoptotic activity varied markedly with both ligand concentration and ligand variant type. To quantify receptor–ligand stoichiometry at the cell membrane, we combined quantitative Stimulated Emission Depletion (qSTED) nanoscopy, simulations, and biochemical assays. Across conditions, signaling complexes consisted of small oligomeric assemblies, with up to three CD95 receptors engaging a trimeric ligand. Despite similar stoichiometry, biochemical measurements revealed substantial differences in ligand oligomerization and binding affinity, with IZ-CD95L exhibiting markedly higher affinity than FLAG-CD95L (*K*_*D*_ of 0.81 nM and 18.4 nM, respectively). Dimeric/trimeric CD95 formed by CD95L complexation or bridged CD95 by mAb enabled flexible intracellular FADD linkage for enhanced signaling.These results indicate that apoptotic potency is governed by the proximity of CD95 receptors, achieved through receptor bridging as well as by a stabilized trimeric ligand, both of which enhance receptor recruitment and binding avidity. Our findings provide mechanistic insight into CD95 signaling and suggest strategies for optimized apoptosis-inducing therapeutics.

**Significance Statement:** The CD95/Fas receptor–ligand system is a central regulator of programmed cell death, yet the earliest molecular events of CD95 activation remain unclear. We systematically compared different CD95L/FasL variants and show that ligand architecture critically shapes receptor engagement and death signaling. An engineered FasL with an isoleucine zipper exhibited larger trimeric ligand fractions, stability, and receptor binding, resulting in stronger apoptotic activity. Super-resolution imaging and simulations indicate that CD95 is mainly monomeric at the membrane and forms small oligomers after ligand binding. The flexibility of intracellular FADD linkage trigerred by dimeric/trimeric or bridged CD95 enables more intense signaling. These findings clarify how ligand structure governs CD95 activation and provide design principles for death receptor–targeted therapeutics.

---

Homopolymerization and cluster formation of cellular membrane receptors is increasingly recognized as an essential facet of cell signaling as well as modulator of physiological responses. Yet, only few techniques are capable of determining the receptor oligomerization states on the cell membrane. In addition, only few studies exist, where different, precise stimuli for apoptosis signal initiation are applied. To better understand the mechanisms and effects of the molecular interactions on the signal initiation and outcome, we employed quantitative stimulated emission depletion (qSTED) nanoscopy combined with biochemical techniques to investigate the stoichiometry of cluster of differentiation 95 CD95/CD95L (Fas/FasL) system. The tumor necrosis factor ligand and receptor superfamily TNFRSF/TNFSF have been widely studied due to their great clinical potential in cancer treatment and immunotherapy. The TNFRSF superfamily consists of 29 members and the TNFSF of 19 members, among which the death receptors (DRs) include cluster of differentiation 95 (CD95), the tumor necrosis factor (TNF), and the tumor necrosis factor related apoptosis inducing ligand (TRAIL). DRs are of great importance, since they trigger the programmed cell death of apoptosis (1). CD95 (Apo 1, Fas or TNFRSF6), a type I transmembrane protein, encompasses three cysteine-rich domains (CRDs) (see Fig.1B), rendering its elongated rod shape (2). The so-called pre-ligand assembly domain (PLAD) in CRD1 is composed of amino acids 1-66 and can dimerize CD95 in the absence of the ligand. PLAD binding has been shown to be weak with a dissociation constant in the µM range (3), wherefore other domains, such as the transmembrane domain, were suggested to contribute to any self-oligomerization (4). CRDs 2 and 3 were shown to mediate the binding to CD95L (5). Recent works showed that CD95 in its resting state on the membrane exists predominantly as monomers along with a small percentage of anti-parallel/parallel dimers (6). Binding of its sole cognate ligand, CD95L, triggers a conformational change of the receptor, recruitment of adaptor protein fas-associated death domain (FADD), and activation of casepase-8 leading to mitochondria involved (type II) (7–9) or independent (type I) apoptosis (10) (see Fig.1A). Ligand-independent apoptosis can be triggered by CD95 agonist antibody alone (e.g. clone: CH11) by placing CD95 in the vicinity for self-activation (11–13), which was assumed to be prevented by inactivation via the PLAD binding site (14). Additionally, agonists play a synergistic role when co-administered with ligands, with a promising example being AMG 655 in cooperation with TRAIL (15). CD95/CD95L system leads to a paradoxical apoptosis or proliferation pathway towards tumorogenesis (16, 17). The latter involves MAPK/NF*κ*B/PI3K pathways (18–21).

**Fig. 1.**
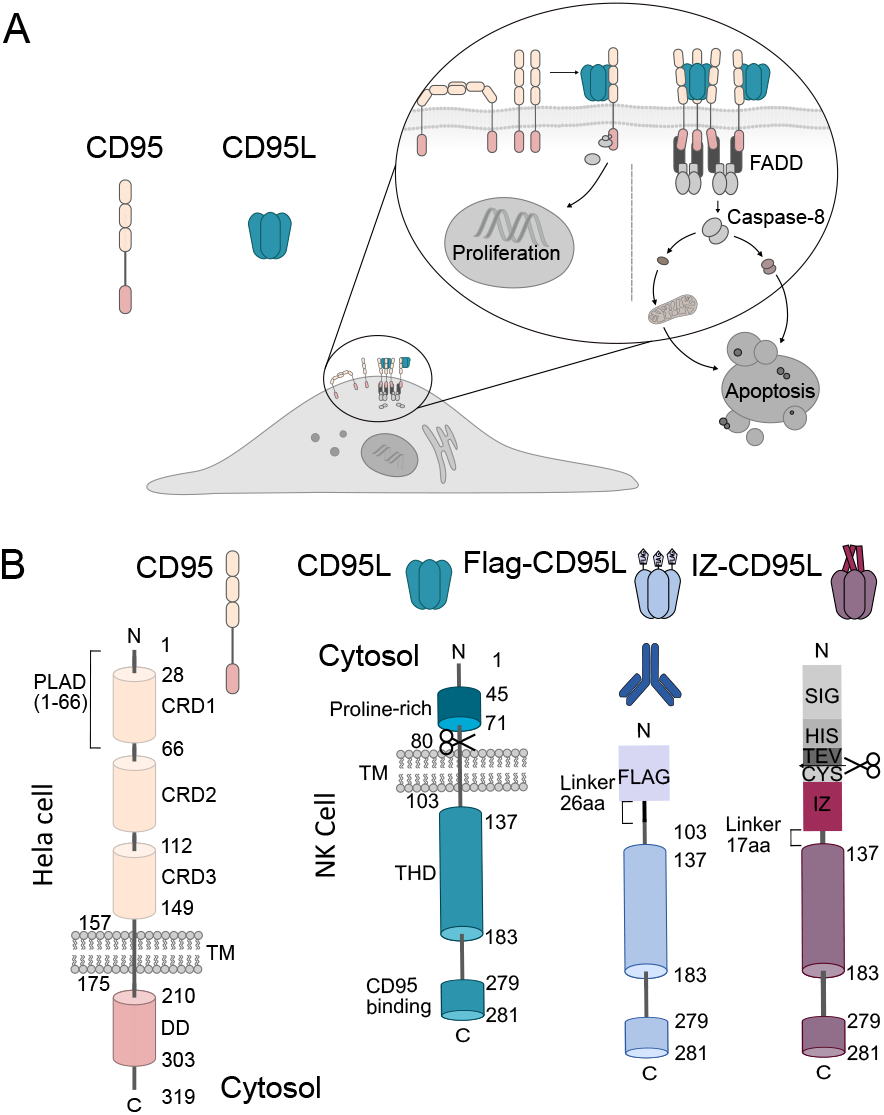
The CD95/CD95L (Fas/FasL) signaling pathway and CD95L variants. A: Illustration of the Cluster of Differentiation 95 (CD95) signaling pathways resulting in proliferation or apoptosis. Here, we investigate the latter, where recruitment of FADD and pro-caspase-8 leads to the crosslinking of intracellular receptor domains. Subsequent downstream signaling evolves via activated Caspase-8 and mitochondria-involved (so-called type II) or -independent (type I) apoptosis signaling, depending on the cell type. B: Sketch of the CD95 receptor and of the CD95 ligand (CD95L) variants. The variants resemble the membrane cleaved CD95L: (i) either the native analog was modified with a *∼*1 kDa Flag affinity tag (Flag-CD95L), (ii) Flag-CD95L was cross-linked with an anti-Flag-tag monoclonal antibody (indicated by the dark blue antibody), (iii) or CD95L was fused with an IZ for trimeric stabilization. For the latter, for correct expression, secretion and purification, additional N-terminal modifications (an extra cysteine for bioorthogonal modification, a TEV cleavage site, a His-tag, and an IL-2 signaling peptide) were added, for the production of ligand protein and detailed description see (22). CRD: cysteine-rich domain; PLAD: pre-ligand assembly domain; TM: transmembrane domain; DD: death domain; THD: TNF homology domain; IZ: isoleucine zipper; CYS: cysteine; TEV(tobacco etch virus): TEV protease recognition sequence; HIS: 8x His-tag; SIG: IL-2 signaling peptide; FLAG: Flag-tag.

CD95L/FasL is a type II transmembrane protein, mainly expressed in immune cells (23). It is characterized by a CD95 binding domain, a TNF homology domain (THD), a transmembrane domain (TM), and an N-terminal proline-rich domain. THD is crucial for homotrimerization, rendering the ligand a bell shape due to the *β* sheet motif in the THD domain. CD95L can be in a membrane-bound or soluble form by metalloproteases cleavage (24), the latter of which shows much less efficiency in apoptosis signaling initiation (25) and was found to trigger non-apoptotic signaling pathways (26). As a shared feature, membrane-bound TNF ligand shows much higher efficiency in signaling transduction than their soluble counterparts. Many efforts have been made to enhance the cytotoxicity of soluble CD95L by increasing structural stability or degree of oligomerization such as fusion with trimerization domains (11, 27–29) or crosslinking ligands with antibodies to enhance local concentration (27). Second, covalent fusion of trimer via a single peptide with single chain antibody (scFv) enhanced targeted antitumoral activity (30). The orientation as well as nanoscale arrangement of CD95L was shown to be important in signaling initiation (31, 32).

The molecular basis of CD95/CD95L (Fas/FasL) binding and nanoscale death inducing signaling complex (DISC) assembly is not yet well understood due to the lack of structural information (33, 34). Understanding CD95/CD95L stoichiometry is crucial for cancer immunotherapy development, yet current techniques lack adequate resolution and specificity. qSTED nanoscopy has been recently used in structural imaging both *in vitro* and *in vivo* due to its sub-diffraction resolution (35–37). However, quantitative analysis of molecular stoichiometry or clustering analysis has not been established yet. In our recent study (6), we developed a quantitative qSTED imaging analysis workflow with molecular sensitivity to aid interpretation of protein stoichiometry and protein-protein interactions. qSTED nanoscopy is based on laser engineering of a depletion laser with a donut shape. Spatial and temporal superimposition of the modulated depletion laser with the excitation laser results in a resolution enhancement by sharpening the point spread function of confocal beam (around 200 nm) to around 40 nm in our case, described by a modified Abbe equation (38):

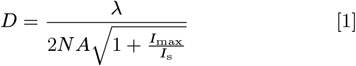

with *λ* being the wavelength of excitation light, *NA* the numerical aperture of objective lens, *I*_*max*_ the maximum qSTED laser intensity and *I*_*s*_ the saturation intensity. To achieve pronounced resolution enhancement, *I*_*max*_ ≫ *I*_*s*_. Furthermore, qSTED resolution can be increased by applying a timing filter, referred to as time-gated qSTED, which was originally developed to improve the resolution of a continuous wave laser (39). In our case, we apply the timing filter on a pulsed qSTED laser with high repetition rate (40 MHz) and high power (1.5 W). So that we imaged single molecule with a mean sigma of 35 nm. Here, we compared apoptosis initiation efficiencies of lab-purified biotinylated IZ-CD95L and commercially available soluble Flag-CD95L (Enzo Life Science), as well as its high-order oligomer by anti-Flag mAb cross-linking (mAb-Flag-CD95L). By employing quantitative qSTED analysis, we reveal unliganded membrane CD95 as monomers form up to trimer after ligand activation (6). We further compared the qSTED spots distribution of the ligand variants, where no nanoscale differences in molecular arrangement were observed. The difference observed in signaling initiation was then proved to be the result of different binding affinities of ligands investigated by surface plasmon resonance (SPR) and blue native PAGE (BN-PAGE) analysis. We conclude that the extent of CD95 apoptosis signaling correlates with ligand affinity.

## Results

### Apoptosis signaling depends on the concentration as well as stoichiometric presentation of the ligand

First, we investigated the apoptosis signal initiation efficiency of the Fas/CD95 ligand variants. Here, we incubated HeLa wild type (WT) cells with the ligands and recorded time-lapse phase contrast micrographs over 20 h. Successful cell apoptosis signaling was detected by cell shrinkage and blebbing. Fig.2A shows snapshots of HeLa WT cells after 2 h, 6 h, and 20 h of ligand incubation at ligand variant concentrations of 28.5 µM. When cells were conditioned with 28.5 µM Flag-CD95L and cross-linked with anti-Flag mAb (100x concentration of Flag-CD95L), first apoptosis events were detected already after 1-2 h. 6 h after ligand incubation, the majority of cells had undergone apoptosis and all signal initiation events were completed. In case of IZ-CD95L, fewer cells died after 6 h, yet whole colonies of cells were apoptotic after 20 h, similarly to the mAb-Flag-CD95L. In case of Flag-CD95L treated cells, an overall low rate of apoptotic cells was observed, with cells being apoptotic mostly some 5-10 h after ligand incubation. The death time of each apoptotic cell was noted and from all apoptosis events over time a sigmoidal curve was obtained, representing the dynamics of cell apoptosis signaling and the eventual cell fate (see Fig.2B). For each experimental condition, an average curve and its standard deviation was calculated and fitted with the Hill equation 3 (see Methods Time-Lapse Imaging). From the fit, the maximum apoptosis percentage [%] and the half life time *t*_*half*_ were derived and plotted in a heatmap (see Fig.2C). In a first comparison, when cells were treated with one of the (IZ-CD95L, Flag-CD95L, or mAb-Flag-CD95L) X-CD95L variants at 5.7 µM, no distinguishable difference between Flag-CD95L and biotin-IZ-CD95L was observed with regard to the total death percentage [%] (27 % and 30 %) and *t*_*half*_ (4.7 h and 5 h). However, mAb-Flag-CD95L exhibited a twofold higher apoptotic percentage of 63 % along with a shorter *t*_*half*_ of around 3 h (Fig.2B). In a second step, the sensitivity of apoptosis signal initiation was checked as a function of ligand concentration. Here, biotin-IZ-CD95L was incubated at 0.57 µM, 5.7 µM, and 57 µM. The bottom graph in Fig.2B shows the result, where 0.57 µM yields negligible apoptosis initiation, identical to a ligand free control experiment. At 5.7 µM ∼ 30 % of apoptotic cells were covered, whereas at 57 µM a threefold enhancement with 90 % apoptotic cells was obtained. This ligand concentration range appears to be necessary to cover the whole range from negligible to near complete apoptosis signal initiation in the exposed cells. Hence, there is a high ligand concentration dependency to initiate the apoptosis signaling pathway along with a ligand variant dependency. To systematically show how the ligand structure and its concentration affect the signal initiation, we extended this experiment to all possible combinations and derived colormaps of the maximum apoptosis percentage [%] and *t*_*half*_ [h], as depicted in Fig.2C. To exclude that in case of IZ-CD95L the biotinylation alters the ligand variant characteristics, also the His-tagged IZ-CD95L was used. In all cases, cell apoptosis was enhanced (*i*.*e*. arising faster and leveling off at a higher percentage), when a higher ligand concentration was used. At 0.57 µM hardly any difference between the ligand variants is detectable, in support of the previous observation that a minimal number of X-CD95L is necessary to trigger the signaling pathway. At 5.7 µM the mAb-Flag-CD95L results in enhanced apoptosis compared to the other cases, whereas at 28.5 µM, IZ-CD95L exhibits equally high apoptosis statistics as the mAb-Flag-CD95L variant, with no detectable differences between biotin-IZ-CD95L and His-IZ-CD95L. The apoptosis statistics of Flag-CD95L, on the other hand, increases only little. At 57 µM a small rise in the apoptosis cases is observed for all variants, along with signatures of saturation. The measurement of mAb-Flag-CD95L at 57 µM was left out, due to the limited availability of the anti-Flag tag antibody. The box plot of Fig.2C summarizes the ligand efficiencies in terms of the apoptosis rate, where the median number of apoptotic cells per hour amount to 0.28, 0.23, and 0.06 [h^−1^] for mAb-Flag-CD95L, biotinylated IZ-CD95L, and Flag-CD95L, respectively. Hence, IZ-CD95L is a highly potent inducer of cell apoptosis signaling in comparison to Flag-CD95L, unless Flag-CD95L is cross-linked with an antibody (mAb-Flag-CD95L). In summary, next to the X-CD95L concentration, its structure and stoichiometric presentation are decisive factors for triggering cell apoptosis. To further probe the origin of these differences, we performed biochemical analyses of the X-CD95L molecular properties.

**Fig. 2.**
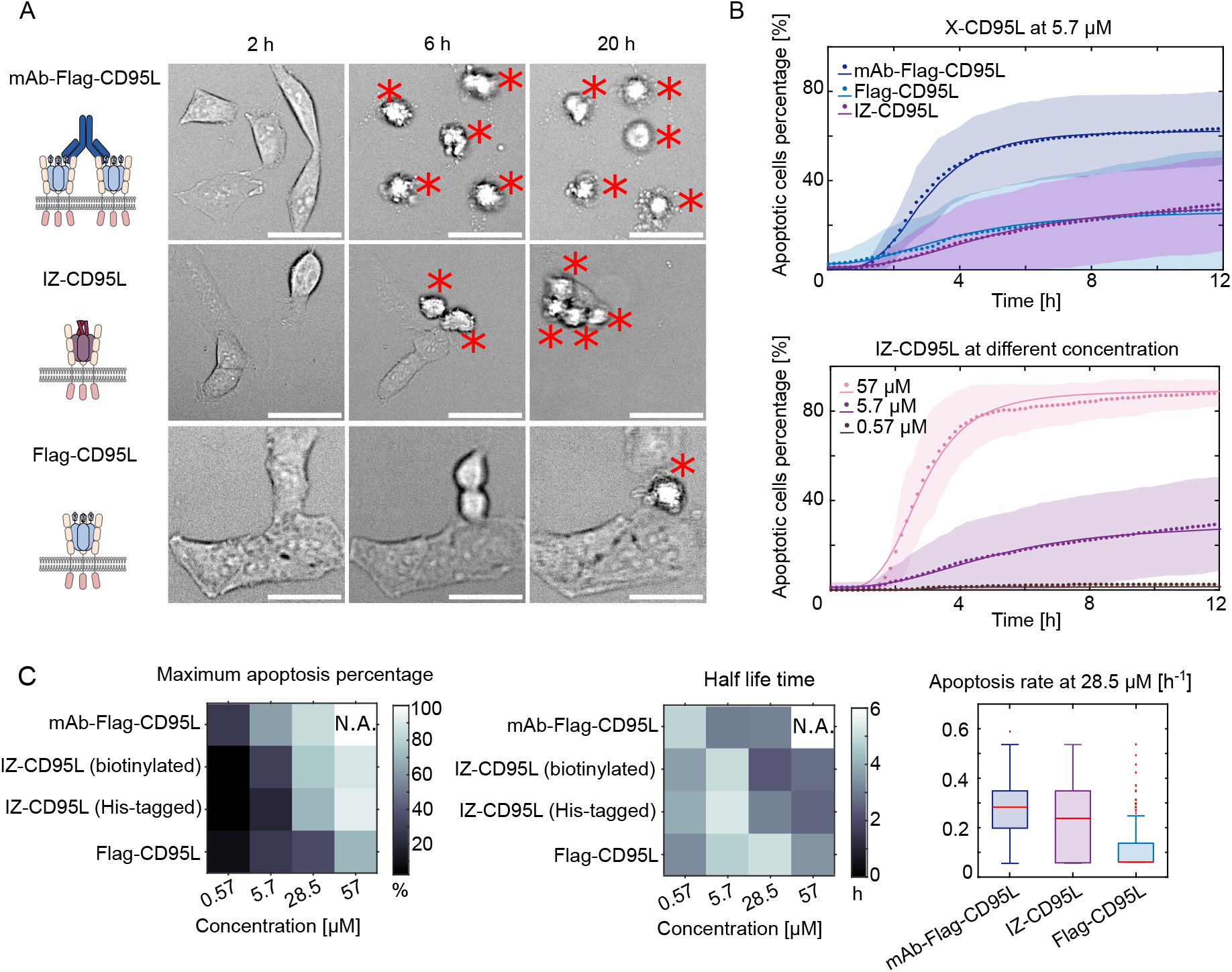
Initiation of apoptosis signaling depends on ligand presentation and concentration. A: Snapshots of cells conditioned with biotinylated IZ-CD95L, Flag-CD95L, and mAb-Flag-CD95L (Flag-CD95L + 100x anti-Flag mAb) ligand variants at 28.5 µM, during time-lapse measurements 2 h, 6 h, and 20 h after ligand addition. Apoptotic cells are marked with an asterisk. Scale bar: 50 µm. B: Apoptosis dynamics curve fitted with Hill equation (see Methods Time-Lapse Imaging for details), to determine the maximum apoptosis percentage, *P*_max_ [%], apoptosis rate, *n*, and half lifetime, *t*_half_ [h]. Upper panel: biotinylated IZ-CD95L (*n* = 2.6, *P*_max_ = 29.9 %, *t*_half_ = 5 h), Flag-CD95L (*n* = 2.2, *P*_max_ = 27.1 %, *t*_half_ = 3.8 h) and mAb-Flag-CD95L (*n*= 3.4, *P*_max_ = 62.8 %, *t*_half_ = 2.8 h) were incubated at 5.7 µM, respectively. Lower panel: biotinylated IZ-CD95 incubated at 0.57 µM (*n* = 3.2, *P*_max_ = 2.6 %, *t*_half_ = 3.7 h), 5.7 µM (*n* = 2.6, *P*_max_ = 29.9 %, *t*_half_ = 5 h), and 57 µM (*n* = 3.8, *P*_max_ = 89.2 %, *t*_half_ = 2.8 h) concentration. Data, Hill equation fits, and standard deviations are presented as dotted lines, solid lines, and shaded areas, respectively. C: Colormaps of maximum apoptosis percentage, *P*_max_ [%], and half lifetime, *t*_half_ [h], of cells treated with CD95L variants at different concentrations. N.A.: a measurement at 57 µM could not be performed. Apoptosis rates were calculated by taking the reciprocal of the death time of each cell: 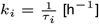. The median values amount to 0.28, 0.23, and 0.06 [h^−1^] for mAb-Flag-CD95L, biotinylated IZ-CD95L, and Flag-CD95L, respectively. Few outliers exceeding the apoptosis rate of 0.7 [h^−1^] are not shown. N > 200 cells per condition.

**Fig. 3.**
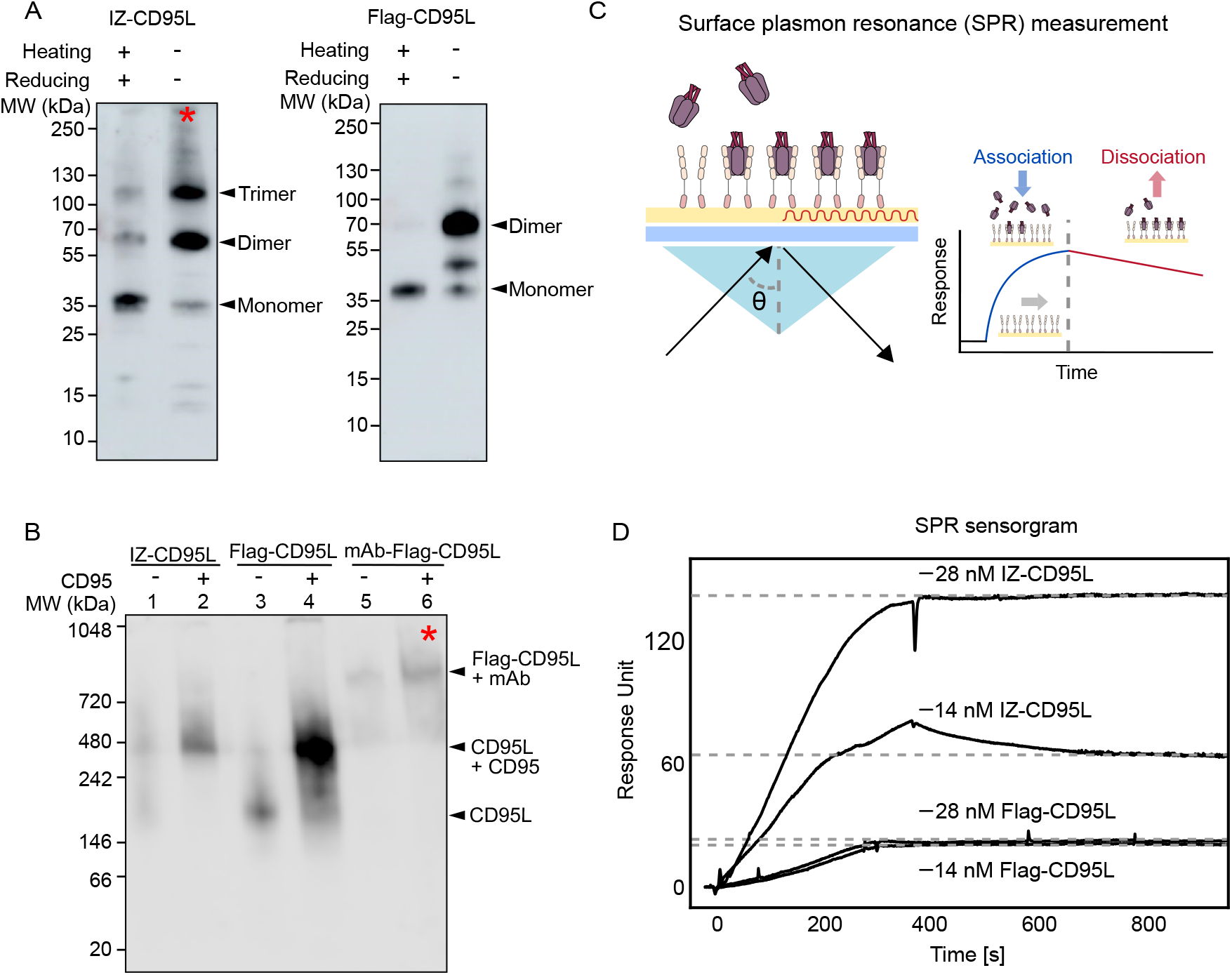
IZ-CD95L trimer stability and CD95 binding affinity is higher compared to Flag-CD95L. A: Ligand stability of IZ-CD95L and Flag-CD95L probed by Western Blot under reducing or non-reducing conditions. Monomers, dimers, or trimers are indicated by black arrows. High molecular weight aggregation is indicated by a red asterisk. B: X-CD95L binding to the CD95 extracellular domain (ECD) probed by Blue Native-PAGE under reducing or non-reducing conditions. IZ-CD95L exhibits a higher fraction of stable trimers and binds more efficiently to CD95 than Flag-CD95L. Only mAb crosslinking of Flag-CD95L results in the formation of higher order complexes. C: Illustration of surface plasmon resonance (SPR) measurements: first, purified ECD-CD95 was immobilized on the sensor chip; second, during the association phase, ligands of a particular concentration were flushed over the chip; third, during the dissociation phase, the sensor chip was flushed with buffers. The resonance angle (*θ*) shifts in response to refractive index changes at the sensor surface, allowing real-time monitoring of binding interactions. Measurements of X-CD95L association and dissociation with ECD-CD95 are plotted in D. D: SPR sensorgrams with equilibrium level indicated by dashed lines. Equation Eq. (2) is used to determine the binding affinities *K*_D_ of X-CD95L to CD95. The *K*_D_ values amounted to 18.4 nM (Flag-CD95L) and 0.81 nM (IZ-CD95L).

### IZ-CD95L is structurally more stable than Flag-CD95L and gives rise to a lower dissociation constant *K*_D_ when binding to CD95

Fig.3A shows the SDS-PAGE result of IZ-CD95L and Flag-CD95L under reducing or non-reducing conditions. For reducing conditions, both Flag-CD95L and IZ-CD95L showed signal bands at ∼35 kDa, corresponding to the expected monomer size with glycosylation. In addition, in case of IZ-CD95L signal bands corresponding to SDS-stable dimers and trimers are detected even after the heating and reducing steps. In case of Flag-CD95L these signal bands did not appear (or only as faint dimer signature), indicating a higher stability of the IZ-CD95L dimer/trimer. Flag-CD95L further exhibited an additional signal band around 45 kDa, corresponding to a fragmented dimer. When identical experiments using non-reducing conditions were performed, signal bands of dimers at ∼70 kDa were detected for both IZ-CD95L and Flag-CD95L. A signal corresponding to the trimer at ∼110 kDa prominently appeared in case of IZ-CD95L and was very dim in case of Flag-CD95L. Based on a rough densitometric estimate from the cropped SDS-PAGE images, the non-reduced IZ-CD95L lane contains approximately 31 % trimer, 42 % dimer, and 9 % monomer. In the non-reduced Flag-CD95L lane, the labeled bands correspond to approximately 57 % dimer and 15 % monomer when normalized to the two major bands only. At the top of this IZ-CD95L lane, a smeared signal appeared (indicated by a red asterisk), which was attributed to higher-order oligomers of IZ-CD95L. In line with previous reports (40), this might arise from domain swapping effects between the IZ of CD95L trimers. To then compare the binding of ligand variants before and after complexation with the extra cellular domain of CD95 (ECD-CD95) proteins, we performed blue native PAGE (BN-PAGE) analyses to avoid possible interference effects of the SDS molecule. In the absence of CD95, trimeric IZ-CD95L (Fig.3B lane 1) and Flag-CD95L (lane 3) migrated to ∼180 kDa. This is larger than the SDS-PAGE result of 110 kDa (see Fig.3A) and is in case of the BN-PAGE typically caused by the larger hydrodynamic size. As for the SDS-PAGE, higher-order oligomers - here at around 480-720 kDa - are observed for IZ-CD95L, which are absent in case of Flag-CD95L. After binding to ECD-CD95, both IZ-CD95L and Flag-CD95L exhibited bands migrating to a size of ∼480 kDa (Fig.3B, lane 2 and lane 4), corresponding to the complexation of three CD95 (∼300 kDa) with X-CD95L (∼180 kDa). Overall, the IZ-CD95L signal is less smeared compared to Flag-CD95L + CD95, indicating an efficient ligand-receptor binding and homogeneous oligomer formation. For comparison, Fig.3B lane 5 and 6 show the antibody crosslinked mAb-Flag-CD95L in the absence and presence of CD95. Here, higher molecular weight signal bands forming between ∼480-900 kDa verify the successful crosslinking of 2 ligands for the high molecular weight fraction around 900 kDa. By comparison with the other lanes, the faint signal band at 480 kDa can be attributed to the uncrosslinked ligand. The presence of CD95 in lane 6 yields slightly larger molecular complexes (red asterisk in Fig.3B), as expected. Next, we probed the binding affinity of X-CD95L to CD95 by surface plasmon resonance (SPR), as it is a highly sensitive, label-free technique, to study scarce materials. Fig.3C shows a sketch of the measurement principle, where the receptor is first immobilized on the sensor chip and thereafter incubated with the ligand (association phase). The sensor is then flushed with buffer, resulting in partial unbinding of the ligand (41). Due to the molecular absorbance on the chip surface, the effective refractive index changes along with the resonance angle of light, that is reflected at the substrate. The resonance angle change is plotted in response units (see Fig.3D) and SPR sensorgrams are recorded for 2 different sample concentrations. Once the equilibrium state (*R*_eq_) was reached at long time-scales, *K*_D_ values were determined according to (42) using equation 2. Accordingly, the *K*_D_ values amounted to *K*_D_ = 0.81 nM for IZ-CD95L (with 63 RU at 14 nM and 142 RU at 28 nM) and 18.4 nM for Flag-CD95L (with 19 RU at 14 nM and 21 RU at 28 nM). This is in line with previously reported values for other artificial ligands binding to CD95 (43). Thus, any complex formation on the cell surface will give rise to larger binding avidities in case of IZ-CD95/CD95 in comparison to Flag-CD95L/CD95. Overall, our biochemical analyses (SDS-PAGE, BN-PAGE, SPR) show that the oligomer of IZ-CD95L is more stable and trimeric in comparison to Flag-CD95L and hence capable to couple CD95 more firmly to produce homogeneous IZ-CD95/CD95 complexes.

In order to then probe the X-CD95L/CD95 complex formation directly on the cell plasma membrane, we performed superresolution STED microscopy and its quantitative data analysis (qSTED).

### qSTED reveals monomeric Fas/CD95 in the cell plasma membrane which oligomerizes into dimers or trimers after complex formation with the ligand

The qSTED measurement and data analysis principle, as previously established (6), are depicted in Fig.4. Fig.4A shows examples of two channel recordings of CD95L (561 nm excitation) and CD95 (640 nm excitation) in confocal and qSTED mode. Here, the lateral resolution from confocal to STED mode changes with the standard deviation of a Gaussian PSF from *σ* = 112 nm to 47 nm. The molecular localization precision is further enhanced with qSTED, where denoising and time gating (3.2-19.2 ns) is applied to ignore early arriving photons that are not completely depleted by the STED pulse with a resolution of 35 nm(39). In addition, a deconvolved image (see Fig.4B-C) is used to find the mass center with x-y coordinates of each spot. The coordinates are then used for 2D Gaussian fitting of the raw STED image and determination of the spot brightness (i.e. the average number of photons < *N*_ph_ > per pixel) and the spot size (i.e. the Gaussian standard deviation, sigma). qSTED data analysis of experimental and simulation results are shown in Fig.4D, where 2D histograms are presented as contour plots with probability density values shown on the right (0.02-0.10 = 2 − 10%). To obtain insights in how the receptor oligomeric state changes the spot characteristics in the qSTED images, we followed the previously established approach (6) to simulate images with monomeric, dimeric, or trimeric CD95 (see Fig.4D, left column). To this end, regions-of-interest (ROIs) containing one, two, or three overlapping 2D Gaussian spots, representing one receptor each, were randomly placed on the image plane. Further, experimentally estimated noise values affecting the sigma and amplitude value were introduced (see Methods qSTED Imaging for details). Fig.4D, right column, shows the result of experimental qSTED data of HeLa WT cells treated with biotin-IZ-CD95L at various concentrations for 2h before fixation. This time frame was chosen as it provides the highest probability to observe CD95 clustering stimulated by ligands: On the one hand, the binding of soluble CD95L to the membrane was reported to saturate within the first 30 minutes after ligand addition (14). On the other hand, first events of apoptosis are observed after 2h (see Fig.2). In Fig.4D a IZ-CD95L concentration dependency of CD95 oligomerization can be observed, where the sigma distribution exhibits a median of ∼4.5 pixels at 5.7 µM, of 5 pixels at 28.5 µM and of 6 pixels at 57 µM. The brightness of spots < *N*_ph_ > per pixel at 5.7 µM and 28.5 µM exhibit a median of ∼3.5 and of ∼7, respectively. At 57 µM the median brightness ∼8. According to the simulation, monomeric CD95 shows a median sigma of ∼4 pixels, whereas dimeric or trimeric CD95 shows a median sigma of ∼5 pixels. The medians of the brightness for simulated monomeric, dimeric, or trimeric CD95 are ∼3, 7, and 10, respectively. We hence associate the condition of 5.7 µM IZ-CD95L predominantly with monomeric CD95 as well as a small portion of CD95 dimers. At 28.5 µM IZ-CD95L a monomeric to dimeric CD95 population is found, whereas at 57 µM IZ-CD95L CD95 forms primarily dimers to trimers. No configurations larger than trimers were measured. These results are in high agreement with CD95 oligomer states reported on HeLa cells before(6), where CD95 after ligand addition formed dimers or trimers, but no higher oligomers. Next, we probed the dependency of the ligand type on the CD95/CD95L oligomerization. To this end, cells were incubated with IZ-CD95L, Flag-CD95L, or mAb-Flag-CD95L at 5.7 µM concentration for 2h before fixation and one control measurement without ligand addition was performed. Fig.5 shows that sigma values of CD95 on the plasma membrane without ligand addition have a median of ∼4 pixels, which was slightly shifted to ∼4.5 pixels after ligand addition in all cases. The brightness distribution was also similar for all cases, with a median value of ∼2.5. CD95 with mAb-Flag-CD95L exhibited a slightly narrower brightness distribution in comparison to Flag-CD95L. This can be caused by the limited accessibility of the CD95 epitope for anti-CD95 staining, when the ligand is already coupled to a large antibody. However, we would expect at least a few events of much higher brightness or sigma, if larger CD95 oligomers existed. Interestingly, neither CD95L variant changes the configuration of CD95 on the membrane at 5.7 µM significantly. By comparison with the without ligand condition it can be estimated that CD95 pre-exists as monomers. This is supported by the monomer/dimer/trimer simulation results, which are overlaid on the without ligand condition. Next to a predominantly monomeric CD95 population, a small fraction of dimers and trimers is also plausible. Note, that absolute sigma and brightness values for 5.7 µM IZ-CD95L are slightly different compared to Fig.4, since here a single antibody staining experiment was performed, whereas in Fig.4 the two channel antibody staining of CD95 and CD95L resulted in a slightly larger intensity offset. In order to cross-check the staining results of CD95, we further analyzed the qSTED measurements with CD95L Antibody-staining (see Fig. S2). Here, for both ligand variants, Flag-CD95L and biotin-IZ-CD95L, sigma values were centered around ∼5-6 [px] and brightnesses < *N*_ph_ > per pixel amounted to ∼2-3 with a slight, yet non-significant upward shift in case of biotin-IZ-CD95L. These small values again indicate a lack of cluster formation. In support of the CD95 data, these data suggest that irrespective of the concentration, no large oligomeric structures evolve and only a recruitment of monomeric to trimeric CD95 to one CD95L occurs.

**Fig. 4.**
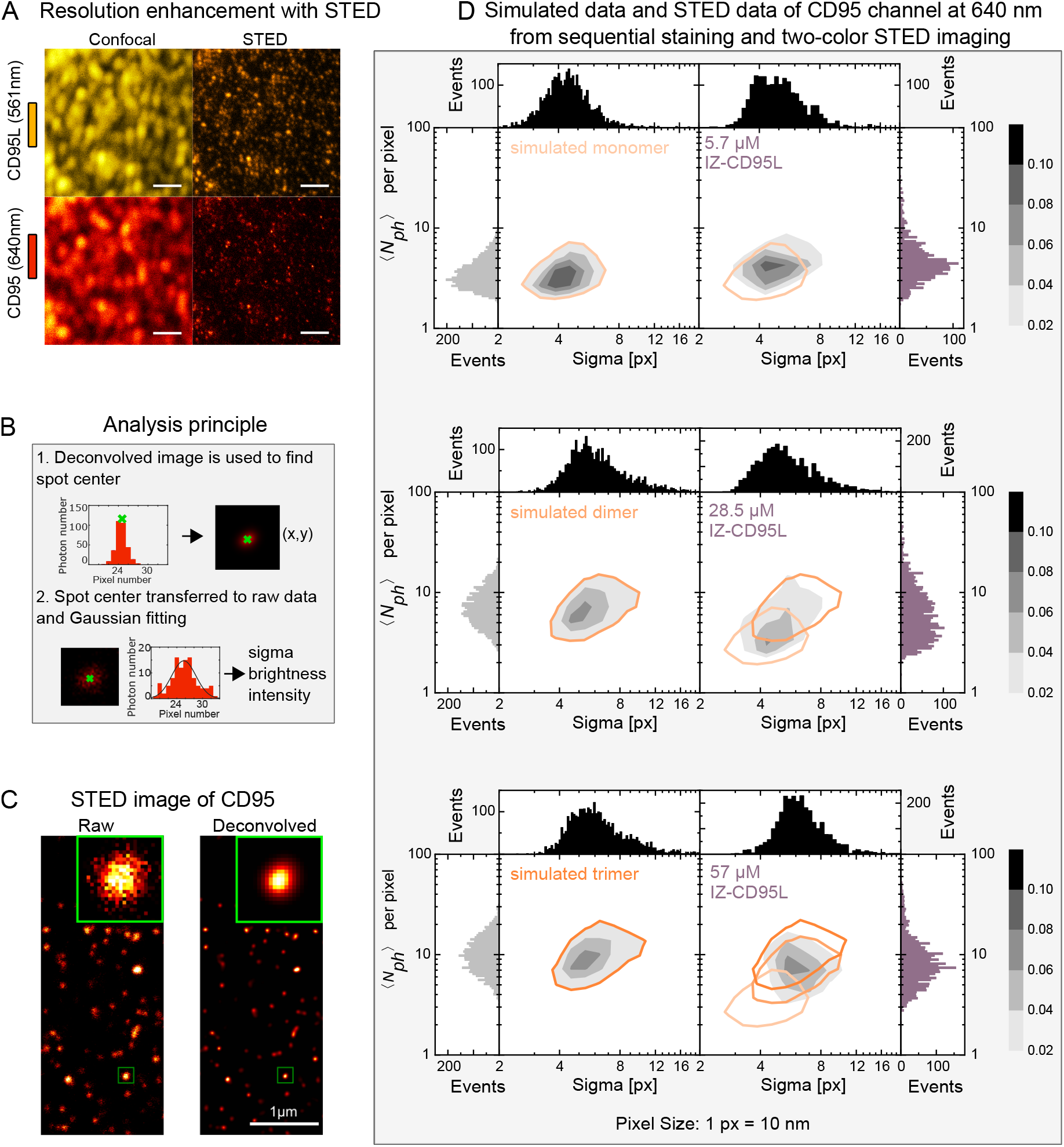
Quantitative STED (qSTED) to probe CD95 oligomerization on the cell membrane by CD95L titration. A: Comparison of two channel data - confocal and STED: CD95L (orange) with excitation at 561 nm and CD95 (red) with excitation at 640 nm alternatively, the images were taken from two color staining experiment. Scale bar: 1 *µ*m. B: Principle of qSTED data analysis: i) Huygens SVI software was used for image deconvolution and to identify spot centers, ii) the coordinates of spot centers were overlaid on raw STED images for Gaussian fittings of the raw data in each ROI. From the fits, the spot brightness (i.e. the average number of photons < *N*_ph_ > per pixel) and the spot size (i.e. the Gaussian standard deviation, sigma [px]) were determined. C: Exemplary STED image (left) of CD95 on the plasma membrane of HeLa WT cell and deconvolved image (right) using Huygens SVI software. D: Two-dimensional probability density representation of brightness (< *N*_ph_ > per pixel) and spot size (sigma [px]) values with log-log scale. Frequency distributions of each parameter are shown on the side and top of each graph. Probability density values are provided as grey colorcode with colorbar given on the right. Left column: simulations of pure monomeric, dimeric, or trimeric CD95 spot distributions. The orange isoline marks the 0.02 = 2% probability density. Right column: measured values for CD95 on the plasma membrane of HeLa WT cells after 2 h incubation with biotinylated IZ-CD95L at 5.7 µM, 28.5 µM, and 57 µM. Number of events are as follows: *N* _monomer_ = 3249, *N* _dimer_ = 3298, *N* _trimer_ = 3316, *N* _IZ-CD95L_ at 5.7 µM = 1810, *N* _IZ-CD95L_ at 28.5 µM = 2862, *N* _IZ-CD95L_ at 57 µM = 2875.

**Fig. 5.**
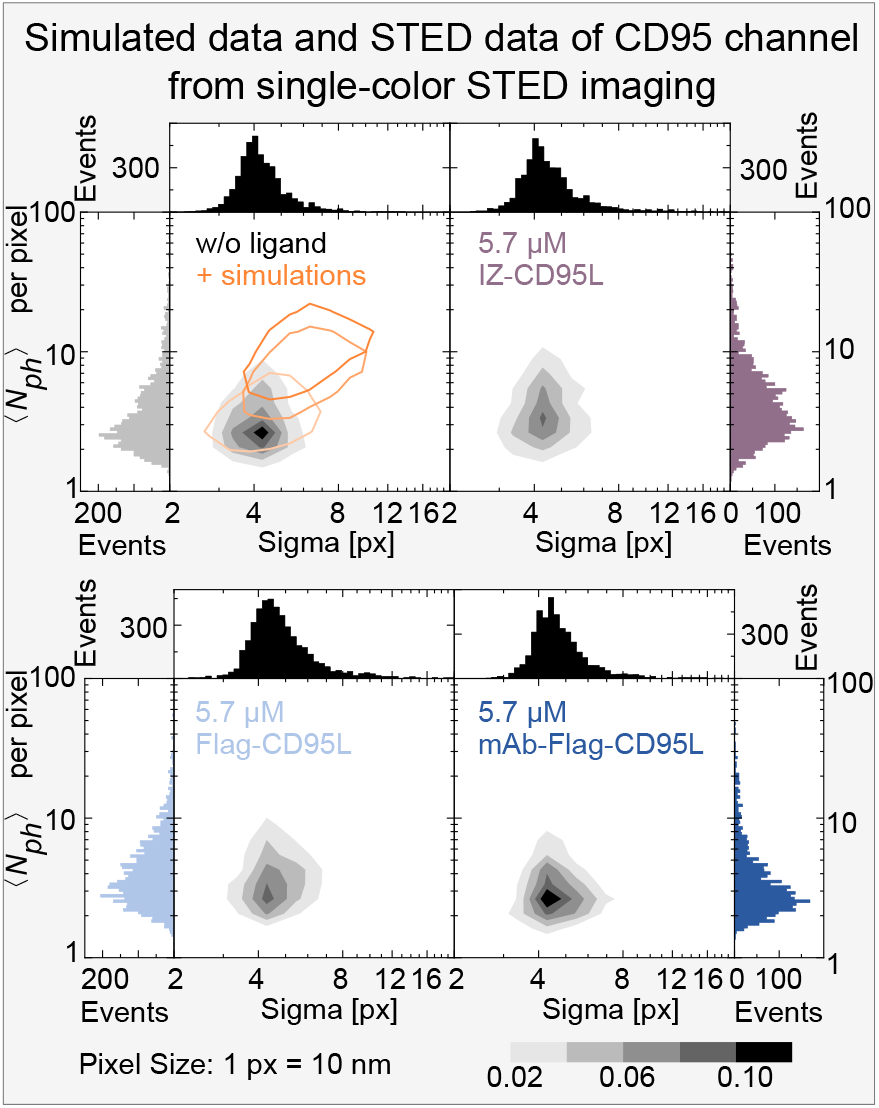
qSTED shows CD95 does not form higher oligomers on the cell membrane upon incubation with CD95L variants. HeLa WT cells were recorded in the absence of CD95L or were treated with 5.7 µM biotinylated IZ-CD95L, Flag-CD95L, or mAb-Flag-CD95L. Two-dimensional probability density representation of brightness (< *N*_ph_ > per pixel) and spot size (sigma [px]) values with log-log scale. Frequency distributions of each parameter are shown on the side and top of each graph. Probability density values are provided as grey colorcode with colorbar given on at the bottom. Orange isolines of simulated monomers, dimers, and trimers were overlaid on the upper left data set where no CD95L was added. Number of events *N* _without ligand_ at 5.7 µM = 3976, *N* _IZ-CD95L_ at 5.7 µM = 4078, *N* _Flag-CD95L_ at 5.7 µM = 4510, *N* _mAb-Flag-CD95L_ at 5.7 µM = 2874.

## Discussion

A direct comparison of the potency of ligand variants was performed using apoptosis dynamics measurements with multi-parameter fitting. We determined the dependency of the apoptosis efficacy and the half life time of the different X-CD95L variants for 0.57-57 µM ligand concentration. Thus, we cover the full range of conditions, with only a few up to all cells undergoing apoptosis. Such approach is further useful as it provides the full dynamic information of the apoptosis signaling efficiency instead of values for selected time points as it is the case in common apoptosis assays (Annexin V, caspase activity evaluation). Previous studies following ligand incubation approaches confirm that ligand concentrations of ∼2.85 µM are necessary to observe substantial apoptosis signal initiation (22, 31, 44). Signaling enhancement was further tested by fusing an IZ motif to CD95L or TRAIL and already demonstrated a high potency to induce apoptotic signaling in HeLa, A549, and HepG2 cells (44). X-CD95L crosslinking via antibodies (e.g. mAb-Flag-CD95L), however, resulted in different outcomes ranging from signaling inhibition (1) to enhanced apoptosis initiation (22). To understand these different responses, it was reasoned that (I) changes in the molecular binding avidity between CD95 and CD95L could play a role and (II) that different oligomerized CD95/CD95L states ranging from small oligomers (monomers to trimers) to large hexagonal arrangements of CD95/CD95L trimer-trimer complexes could develop (1). Thus, we performed biochemical analyses to probe the structural stability of the different ligand variants, their oligomeric states, and, consequently, their binding avidities. It is important to note, that CD95L is generally considered to exist in a trimeric state, albeit several previous studies showed that the soluble form of CD95L is partioned in monomers, dimers, and trimers (22, 45–47). Due to the missing groove region in case of CD95L monomers, the specific finding in this study further excludes the existence of CD95L/CD95 monomer-monomer complexes. The long discussed limited efficacy of soluble CD95L vs. membrane bound CD95L (16, 31), may hence be a direct consequence of the higher amount of trimeric and structurally stabilized CD95L on the cell plasma membrane in comparison to the secreted or cleaved and purified soluble CD95L. For both ligand variants, Flag-CD95L and IZ-CD95L, substantial monomeric and dimeric fractions are detected for non-reduced SDS-PAGE conditions. While the SDS detergent contributes to the protein denaturation, and may hence exaggerate the fraction of dissociated protein, in our previous study we characterized IZ-CD95L during different protein purification steps with SEC (22) and, from the beginning, found a dominant trimeric fraction (60 %), followed by a non-negligible dimer fraction (30 %). Furthermore, a direct consequence of the apparent reduced stability in case of Flag-CD95L appears to directly translate into a 20 times lower binding avidity *K*_*D*_ (Fig. 3D). Next to these differences in oligomeric fraction, Flag-CD95L may also bind less strongly to CD95-ECD than IZ-CD95L. While the amino acids mediating the receptor-ligand interaction are identical in both cases, the IZ peptide self-assembles into a highly stable, parallel three-stranded *α*-helical coiled coil (48). The subunits of the oligomeric IZ-CD95L could hence be more parallelized and oriented in comparison to Flag-CD95L. The binding interface between CD95 (Fas) and CD95L (Fas ligand) has been well characterized and involves defined structural domains on both molecules. The interaction between CD95 (Fas) and CD95L is primarily mediated by the third cysteine-rich domain (CRD2-3) of CD95 and the TNF homology domain (THD) of CD95L (49). These binding sites are conserved, suggesting that different CD95L constructs, such as Flag-CD95L and IZ-CD95L, engage the same receptor epitope. Therefore, differences in binding avidity are unlikely due to changes in the binding interface, but rather to variations in oligomerization and spatial organization. In particular, the more stable and ordered trimeric structure of IZ-CD95L may promote improved receptor clustering, while less organized forms may introduce steric hindrance and reduce binding efficiency. Still, antibody crosslinking of Flag-CD95L (mAb-Flag-CD95L condition) not only restores, but supercedes the efficiency of IZ-CD95L. We reason that the coupling of the anti-Flagtag antibody induces a higher local concentration of dimeric and trimeric CD95L resulting in CD95 receptors just at the right distance for intracellular crosslinking and downstream signaling. In addition, antibody binding could support the correct orientation and provide a higher structural stability of Flag-CD95L. Finally, different oligomerization models for CD95/CD95L on the membrane have been suggested (1, 6). While our data excludes the existence of large supramolecular complexes (such as the hexagonal trimer-trimer configuration suggested in (1)), here and in our previous study (6), only small oligomers up to CD95/CD95L trimer-trimer states were found. It should be noted that several studies reporting about the large oligomeric complexes (1) used biochemical techniques that required high molecular concentrations (mM regime). While we find a slight concentration dependency with higher oligomers forming at higher molecular concentrations (6), large oligomeric complexes may indeed appear only at high concentrations. The concentration dependency of the signaling efficiency is hence readily explained, as the probability of ligand - receptor complex formation is enhanced when more ligands are presented (50). Direct measurements on the membrane, however, are only possible since the invention of super-resolution approaches and have led to reports of substantially smaller oligomeric states (51).

We obtain insights from different molecular concentrations and using systematically modified ligands. Only X-CD95L/2CD95 or X-CD95L/3CD95 can trigger apoptosis initiation, since the activation of caspase-8 needs at least two pro-caspase-8 in close proximity. FADD recruitment has been reported to depend on the X-CD95L/CD95 complexation on the plasma membrane (14), with more than fourfold better efficiency of FADD recruitment observed after 1 h in case of IZ-CD95L in comparison to sCD95L.

### Proposed model of CD95/CD95L variant binding for apoptosis signal initiation

The reported findings support the following model of CD95/CD95L variant binding (Fig. 6). Apoptotic signaling initiation is strongly dependent on the concentration of X-CD95L. In the absence of ligand, CD95 predominantly exists as monomers at the plasma membrane, although a minor fraction may form small oligomeric assemblies, including previously described inactive dimers. Upon engagement of X-CD95L with a single CD95 molecule, up to two additional CD95 receptors can be recruited to a trimeric X-CD95L, resulting in formation of a signaling-competent complex. In the complex, dimerized CD95 recruit intracelullarly FADD and lead to dimerization/activation of caspases. Flag-CD95L exhibits approximately twentyfold lower affinity for CD95 than IZ-CD95L, as illustrated by the dashed lines and increased spacing between Flag-CD95L and CD95 (upper panel) compared with IZ-CD95L and CD95 (middle panel). Nevertheless, Flag-CD95L can be antibody-crosslinked to generate higher-order assemblies, including dimerized trimers (bottom panel), thereby stabilizing receptor engagement. Although the stronger intrinsic binding of IZ-CD95L promotes apoptotic signal initiation relative to Flag-CD95L, antibody-mediated crosslinking of Flag-CD95L enhances signaling efficiency beyond that achieved by IZ-CD95L alone. No large supramolecular complexes were detected under any condition; oligomerization was restricted to CD95 trimers. The proximity of activated CD95 receptors therefore appears to be the critical determinant for efficient signal initiation. In the presence of antibody-crosslinked Flag-CD95L, this spatial arrangement is optimal for recruitment of the adaptor protein FADD and pro-caspase-8. The latter undergoes concentration-dependent autocatalytic activation, which subsequently drives downstream apoptotic signaling. CD95 molecules crosslinked by monoclonal antibodies (mAbs) are referred to as “bridged CD95” complexes (e.g., (CD95)_2_–(CD95)_2_ or (CD95)_3_–(CD95)_2_), which are structurally distinct from CD95 dimers formed through CD95L binding, as illustrated in the upper and middle panels. The critical intermolecular distance for both dimeric and bridged CD95 is estimated to range from 7 to 10 nm (10, 12). Moreover, the flexible linkage of FADD within the intracellular domain enables highly dynamic interactions, such that FADD assembly within (CD95)_2_–(CD95)_3_ complexes promotes more robust downstream signaling.

**Fig. 6.**
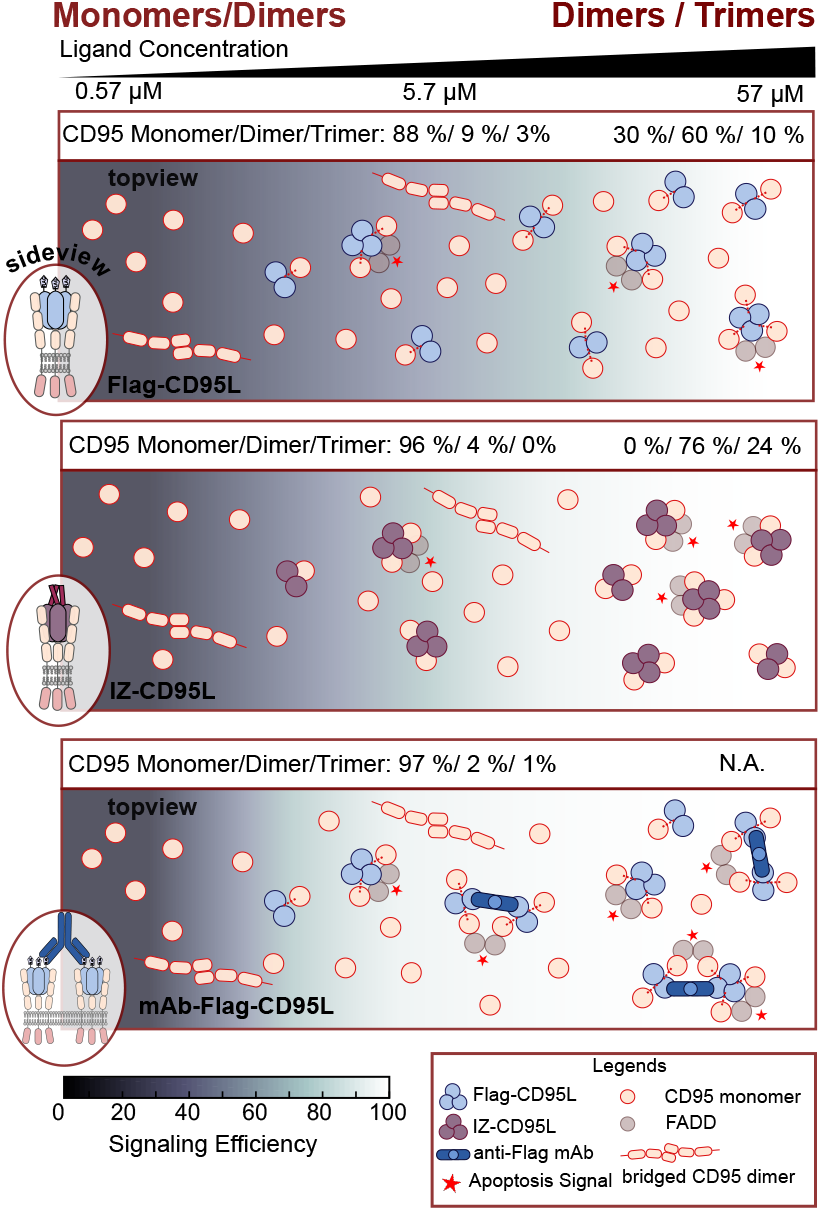
Graphical summary of X-CD95L complexation with CD95 on the cell membrane. From top to bottom: Flag-CD95L, IZ-CD95L, and mAb-Flag-CD95L (Flag-CD95 + 100x anti-Flag mAb). Apoptosis signaling efficiency (black - blue - white color scale) increases with CD95L concentration. Accordingly, in absence of the ligand (left side of the graph) CD95 pre-exists as monomer. It oligomerizes to dimers or trimers, after complexation with the ligand (right side of the graph). A few percent of CD95 per previously shown to form inactive dimers via their PLAD domains. The higher the structural stability and binding affinity of X-CD95L to CD95 as well as ligand concentration, the more X-CD95L/CD95 complexes with CD95 dimers or trimers form. Differences in the X-CD95L binding affinity are presented as loose (red dashed lines for Flag-CD95L in first and last row) or tight binding (shown as a lack of spacing for IZ-CD95L between ligand and receptor). X-CD95L binding brings two CD95 in close proximity, which recruit the adaptor protein FADD and pro-caspase-8. This dimerization/high local concentration triggers the autocatalytic cleavage of matured caspase-8 and the apoptosis downstream signaling (indicated by red asterisks).

## Materials and Methods

### Cell Culture and Sample Preparation

HeLa WT cells (CCL-2™, ATCC, Manassas, VA, USA) were cultured in T25 cell culture flasks (Sarstedt, Nümbrecht, NRW, Germany) with DMEM medium (Gibco, Life Technologies Inc., Carlsbad, CA, USA) containing 10 % FBS and 1 % penicillin/streptomycin (P/S) solution at 37 °C in an incubator humidified with 5 % CO_2_ (v/v). Cells were passaged at *∼*70 % confluency and detached using trypsin-EDTA solution. For qSTED sample preparation, cells were seeded on sterile glass slides in 12 well plates (Sarstedt) with each well containing 100-150x 10^4^ cells. Cells were left to adhere for at least 12 h and were washed with DPBS prior to ligand incubation. Flag-CD95L (Enzo Life Sciences, Farmingdale, NY, USA) with or without enhancer (mouse anti-Flag mAb) or IZ-CD95L (22) were diluted in 300 µl pre-warmed L15 medium containing 10 % FBS and 1 % P/S at the stated concentrations and cells were incubated with them for 2 h at 37 °C in the incubator.

### Time-Lapse Imaging and Apoptosis Dynamics Analysis

Time-lapse measurements were performed on an IX83 microscope (Olympus, Evident, Waltham, MA, USA) and MFIS-SSTR Nikon microscope (A1HD25-NSTORM) using a 20x air objective. Cells were seeded on a 8 well plate (Sarstedt) one day before the measurement with *∼*50 % confluency and mounted on a temperature controlled stage (PeCon, Erbach, BW, Germany) at 37 °C. X-CD95L were diluted in L15 complete medium (10 % FBS and 1v% P/S) and added to the cells *in situ* on the microscope just before the start of the measurement. Time-lapse videos in phase contrast or bright field mode were acquired for each condition using the CellSense Dimensions Software (Evident Scientific, Hamburg, Germany). Snapshots of five positions from each well were taken every 5-15 min overnight with a Z-drift compensation system IX-ZDC. Image analysis was performed using ImageJ (52), where every single apoptotic cell was identified by characteristic morphology changes (e.g. cell blebbing and apoptotic bodies) where the frame number was exported and converted into the death time. The resulting percentage of apoptotic cells *P*(t) was fitted with a Hill equation using MATLAB2018 (53) (Mathworks, MA, USA):

### SDS-PAGE and Western Blot

100 ng IZ-CD95L or Flag-CD95L were reduced and boiled at 95 °C for 5 min in reduced 1x Laemmli buffer (Bio-Rad). For non-reducing condition, samples were treated in 1x Laemmli buffer without heating. They were loaded together with 5 µl pre-stained protein ladder (Thermo Fisher Scientific Inc., Waltham, MA, USA) and vertically separated on a 4-20 % pre-cast gel using Mini-PROTEAN Tetra Cell (Bio-Rad, Hercules, CA, USA). Afterwards, protein bands were transferred to a PVDF membrane by a semi-dry Trans-Blot Turbo Transfer System (Bio-Rad). IZ-CD95L and Flag-CD95L were probed with the primary antibodies mouse anti-His-tag (clone: J099B12, dilution 1:1000) (Biolegend, San Diego, CA, USA) or mouse anti-Flag-tag antibody (ENZO, enhancer, dilution 1:1000) at 4 °C overnight, followed by incubation with horseradish peroxidase (HRP) linked goat-antimouse secondary antibody (Cell Signaling Technology, Danvers, MA, USA, dilution 1:1000) at r.t. for 1 h. After the development with an ECL substrate (Bio-Rad), images were taken with an Amersham™ Imager AI680 (Cytiva, Marlborough, MA, USA).

### Blue Native Page

2.86 pmol IZ-CD95L or Flag-CD95L were incubated with 10x ECD-CD95 (Sino Biological, Beijing, China) or, additionally, with 10x anti-Flag-tag antibody (ENZO) at 37 °C for 2 h. Afterwards, the ligand alone or ligand/receptor complexes were loaded with 10 % glycerol and 0.5 % G250 on a 4-20 % pre-cast gel (Bio-Rad) together with 10 µl unstained protein standard (Invitrogen, Waltham, MA, USA). Electrophoresis was performed at 4 °C/100 V with the cathode buffer (50 mM tris/ 192 mM glycine/ pH 8.3/ 0.02 % G250) and the anode buffer (50 mM tris/192 mM glycine/pH 8.3). Cathode running buffer was exchanged with (50 mM tris/ 192 mM glycine/ pH 8.3) when G250 had migrated through half of the gel. Afterwards, proteins were transferred to a PVDF membrane followed by a Western Blot. The blot was probed with anti-CD95L rabbit antibody (Abcam, Cambridge, UK, ab134401, dilution 1:1000) at 4 °C overnight followed by incubation of goat-anti-rabbit HRP-linked secondary antibody (Cell Signaling Technology, dilution 1:1000) for 1 h at r.t.

### Surface Plasmon Resonance Measurement and Data Analysis

Surface plasmon resonance measurements were performed using the 2-channel system Reichert 2SPR (Ametek Reichert Technologies, Depew, NY, USA). Either a nitrilotriacetic acid (NTA) derivatized linear polycarboxylate hydrogel chip (Xantec bioanalytics, Germany, SCR NiHC30M) or a streptavidin derivatized linear polycarboxylate hydrogel chip (Xantec, SCR SAH30M) were used for CD95 ECD (his-tagged or biotinylated) immobilization. X-CD95L was titrated with different concentrations on the CD95 chip with a flow rate of 30 µl/min for 6 min followed by washing. The chips were regenerated by washing with ethylenediaminetetraacetic acid (EDTA) and recharged with Ni^2+^ or regenerated by 4 M MgCl_2_. *K*_D_ values were estimated from the sensor graphs from the steady state by using equation 2.

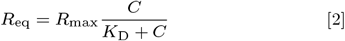

where C is the concentration of the analyte, *R*_eq_ is the equilibrium response unit (RU), *K*_D_ is the equilibrium dissociation constant, and *R*_max_ is the maximum theoretical response unit (RU) to be observed when all available binding sites on the sensor surface are occupied by analyte.

### Immunofluorescence Staining for qSTED Imaging

Considering CD95 and CD95L binding with fast kinetics (14), qSTED staining was performed always after 2 h incubation with ligand. After this time, cells were briefly washed with DPBS and then fixed with 4 % methanol-free formaldehyde (Thermo Fisher) diluted in DPBS at r.t. for 10 min. Afterwards, cells were rigorously washed three times with DPBS for 5 min with shaking. To increase the antibody binding efficiency, samples were treated first with preheated retrieval buffer (5 % Urea (w/v) /100 mM tris/ pH 9.5) at 95 °C for 10 min followed by three times of 5 min washing with DPBS. Next, cells were permeabilized with 0.2 % triton X-100 in DPBS and washed 3x 5 min afterwards. To reduce unspecific staining, cells were passivated with blocking buffer (1 % BSA / 22 mg/ml glycine /0.1 % tween-20 in DPBS) at r.t. for 30 min. For two-color STED, the cells were stained with 10 µg/ml anti-CD95L (Abcam, ab134401, rabbit polyclonal antibody) and 5 µg/ml anti-CD95 (Biotium, Fremont, CA, USA, clone:B-R18, mouse monoclonal antibody) overnight at 4 °C, followed by 1 h incubation at r.t. in the dark with secondary antibodies Abberior STAR RED goat-anti-mouse and Abberior STAR ORANGE goat-anti-rabbit (Abberior) both at 5 µg/ml. Two color staining and STED imaging was performed to validate the system and serves as control. Secondly for single color STED, the cells were only stained with 10 µg/ml CD95 antibody (Invitrogen, clone:JJ0942, rabbit monoclonal antibody) overnight at 4 °C followed by Abberior STAR ORANGE goat-anti-rabbit staining for 1 h at r.t. Cells on cover slips from a 12-well plate were flipped on imaging slides and cured using ProLong™ diamond antifade mountant (Invitrogen) at 4 °C o/n. Successful staining of cells were briefly checked by IX83 microscope (Olympus, Evident, Waltham, MA, USA) or Nikon N-STORM system based on a Ti-E inverted microscope (Nikon, Japan).

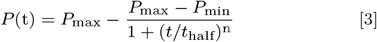

with parameters *P*_min_ and *P*_max_, as the minimal and maximal percentages of apoptotic cells, *t*_half_, as the characteristic time when half of all cells are apoptotic. The Hill coefficient (cooperativity coefficient) *n* indicates how steep *P*(t) rises, i.e. whether the signal in all cells is processed equally fast.

### qSTED Imaging

qSTED imaging was performed on a customized Abberior Expert Line Setup (Abberior Instruments, Göttingen, Germany) (54). All samples were imaged with one or two linearly polarized excitation lasers at 561 nm (2.9 µW) and 640 nm (19 µW) and using a single circularly polarized qSTED depletion laser at 775 nm (31.50 mW). A 100x oil objective (NA 1.4, UPLSAPO 100x Oil) (Olympus, Hamburg, Germany) was used. Before the measurements, excitation and depletion lasers were aligned using 0.1 µm TetraSpeck beads (Invitrogen). First, the confocal mode was used to find the cells, followed by recording 10 ROIs of 5 µm x 5 µm size in qSTED mode. Each ROI was scanned using a 10 nm pixel size, 3.00 µs dwell time, and 5 frames in the Imspector Software (Abberior). Fluorescence was first separated into perpendicular and parallel polarization components using a polarizing beam splitter followed by fluorescence bandpass filter cubes Cy3 (Abberior ET608/ 44) or Cy5 (Abberior ET685/ 70M) for each polarization direction. Detection was achieved by four avalanche photodiodes (APDs, Excelitas, Pittsburg, PA, USA) synchronized with a time correlated single photon counting unit (TCSPC, PicoQuant, Berlin, Germany). While the hardware enables to analyze fluorescence polarization effects, in this work, only the total fluorescence signal (i.e. the sum of the parallel and perpendicular polarization channels) was evaluated. We further identified native expression of mCD95L on HeLa WT cells and we tested passivation of native mCD95L by either anti-CD95L antibody or purified CD95 ECD protein.

### qSTED Data Analysis

The resolution of qSTED images was first enhanced by time-gating the first 3.2-19.2 ns using the self-written LabView 2020-based software AnI-3SF https://www.mpc.hhu.de/software/mfis-2021(LabVIEW 2020, NI, USA). Huygens Professional (HuPro Version 21.10.1p2 64b or later) was used to deconvolve the images, which was performed via the classic maximum likelihood estimation (CMLE) algorithm, with a signal-to noise ratio (SNR) of 4, a maximum of 25 iterations, an acuity of 97, and a relative background fixed to 0.7. After deconvolution, Huygens Object Analyzer was used to determine the mass centers along the X and Y axis (CmassX CmassY) of each spot. Here, watershed was set as the segmentation mode with an absolute threshold of 7 and a fragmentation percentage of 50 %. The garbage volume was set to 1 voxel. Afterwards, the mass center of each spot was overlaid on photon-gated raw images as ROIs (11 x 11 pixels) and fitted with 2D Gaussians using AnI-3SF with a minimal spot size of 10 pixels, a segmentation object threshold of 10, and with a fixed background. Objects touching image borders were ignored. Sigma and brightness values obtained from the fit to each spot were plotted, with the number of photons per pixel (< *N*_ph_ > per pixel, Nph for short) and the standard deviation (*σ*) of each spot were plotted in the form of a 2D histogram using OriginLab (2020 or later, MA, USA) and probability density values indicated.

### qSTED Simulations

The simulation of receptor oligomers recorded with qSTED resolution (with FWHM of 82 nm) was done with LabView 2020 (National Instrument, Austin, TX, USA). The following routine was applied: i) monomers, dimers or trimers were modeled as single, two, or three 2D Gaussian distributions placed at defined coordinates based on estimated intermolecular spacings in a ROI with 25 x 25 pixel size (see Fig. S9A). The sigma and amplitude of each Gaussian was taken from experimental qSTED images of CD95 on HeLa WT membrane without the ligand, where CD95 primarily pre-exists as monomers (the fraction of inactive dimers amounts to only *∼*4%)(6, 14), with Gaussian noises introduced. We specified the simulation parameters of each condition — monomer, dimer, and trimer as: *σ* and *σ*_noise_; gaussian amplitude and amplitude noise; x and y coordinates; background noise; circularity; see Table S4. ii) From the experimental data, the mean and standard deviation of the distributions of *σ* and amplitudes of all spots were obtained following qSTED data analysis and used as Gaussian noises (Table S3). In summary, for monomer spots the final Gaussian width was set to its mean value *σ* = 3.5 px (with 30 % relative noise, *σ*_noise_ = 0.3) and the amplitude was set to its mean value of 2.5 ph (with 100 % amplitude noise). For dimer and trimer spots, the final Gaussian width was set to its mean value *σ* = 4.5 px (and *σ*_noise_ = 0.3) and the amplitude was set to its mean value of (2.5 ph ± 100 %) (< *N*_ph_ > per pixel, ph for short). iii) 1000 of these ROIs were randomly placed on an overview image with 500 x 500 pixel size. ROIs touching the boundaries were excluded (positions constrained to 0–475 px in both axes). Poisson-distributed background noise (mean = 0.295 ph/px) was added to every pixel. The noise level was determined from the average background intensity from CD95 qSTED images acquired under ligand-free conditions. While the initial input parameters produced simulation output values *σ*, intensity, and noise that slightly deviated from the experimental target values, we ran simulations iteratively to fine-tune the input parameters until the simulated distributions closely matched the experimental monomer dataset. The simulated images were saved as U16 TIFFs and analyzed following the qSTED data analysis workflow stated above.

## ACKNOWLEDGMENTS

CM, XS and NB acknowledge financial support by the DFG Collaborative Research Center 1208 “Identity and dynamics of biological membranes” (project ID 267205415). CM and AN acknowledge financial support by the ‘Freigeist-fellowship’ of VolkswagenFoundation as well as the financial support of CM and XS via ‘Momentum’ of VolkswagenFoundation. The authors acknowledge the Deutsche Forschungsgemeinschaft

(DFG, German Research Foundation) and the state of North RhineWestphalia for funding the custom MFIS-SSTR Nikon microscope (A1HD25-NSTORM) within the program Major Research Instrumentation as per Art. 91b GG with the funding ID INST 208/797-1 FUGG to Claus Seidel and CM, and the custom Abberior Instruments Expert Line STED microscope for MFIS-STED within the program Major Research Instrumentation as per Art. 91b GG with the funding ID INST 208/741-1 FUGG to Claus Seidel. We thank Prof. Alexej Kedrov (Synthetic Membranesystems, Heinrich-Heine University Düsseldorf) for providing the Amersham Western Blot Imager and the 2-channel system Reichert 2SPR.

