## Supplementary Information (SI) for "CD95/Fas Apoptosis Signal Initiation Depends on the Ligand Oligomerization State and Formation of Small Ligand-Receptor Complexes"

#### This PDF file includes:

Figs. S1 to S4
Tables S1 to S3

### Supporting Figures

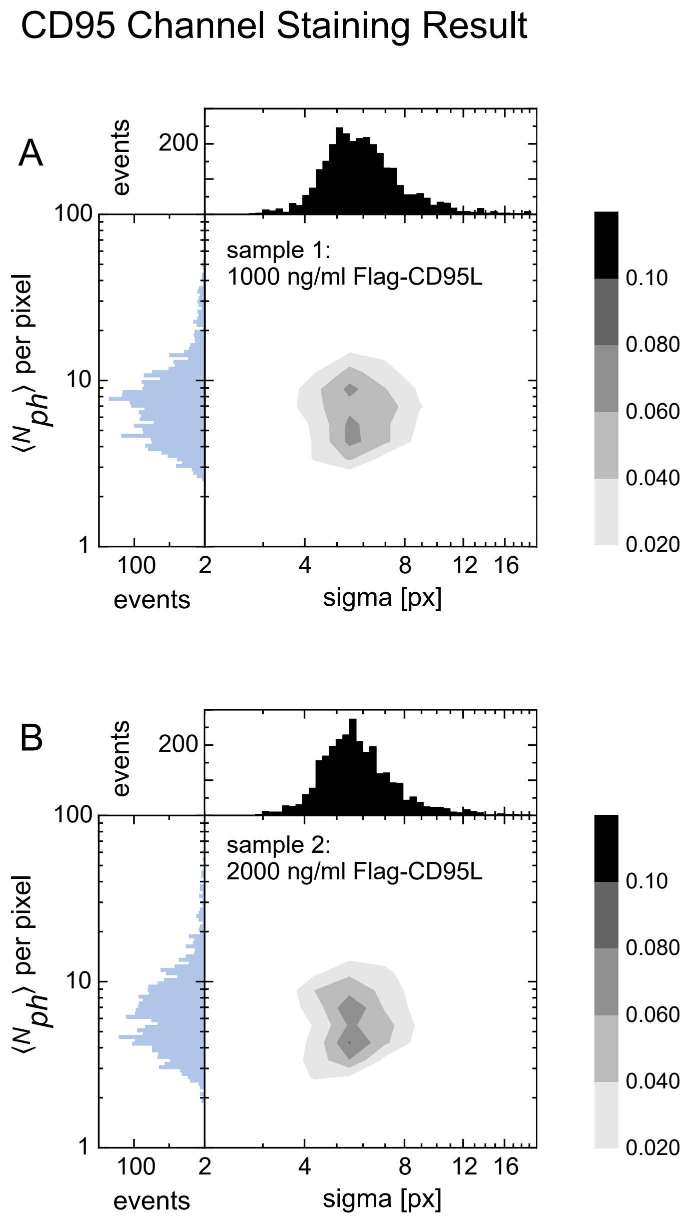


**Fig. S1.** CD95 results from quantitative spot analysis of two color stained STED images. Two-dimensional probability density representation of brightness ($N_{\mathrm{ph}}$ per pixel) and spot size (sigma [px]) values with log-log scale. Frequency distributions of each parameter are shown on the side and top of each graph. Probability density values are provided as grey colorcode with colorbar given on the right. Flag-CD95L was added at 1000 ng/ml and 2000 ng/ml concentrations and incubated on Hela WT cells for 2 h at r.t. After washing and fixation, cells were incubated with primarily 5 µg/ml anti-CD95 (Biotium, mouse monoclonal, clone:B-R 18) and 10 µg/ml anti-CD95L (Abcam, rabbit polyclonal, ab134401) overnight at 4 ℃ followed by washing and incubation of secondary antibodies goat anti-rabbit AbSTAROrange and goat anti-mouse AbSTARRed (both from Abberior at 5 µg/ml) at r.t. for 1 h in the dark.

### Supporting Tables

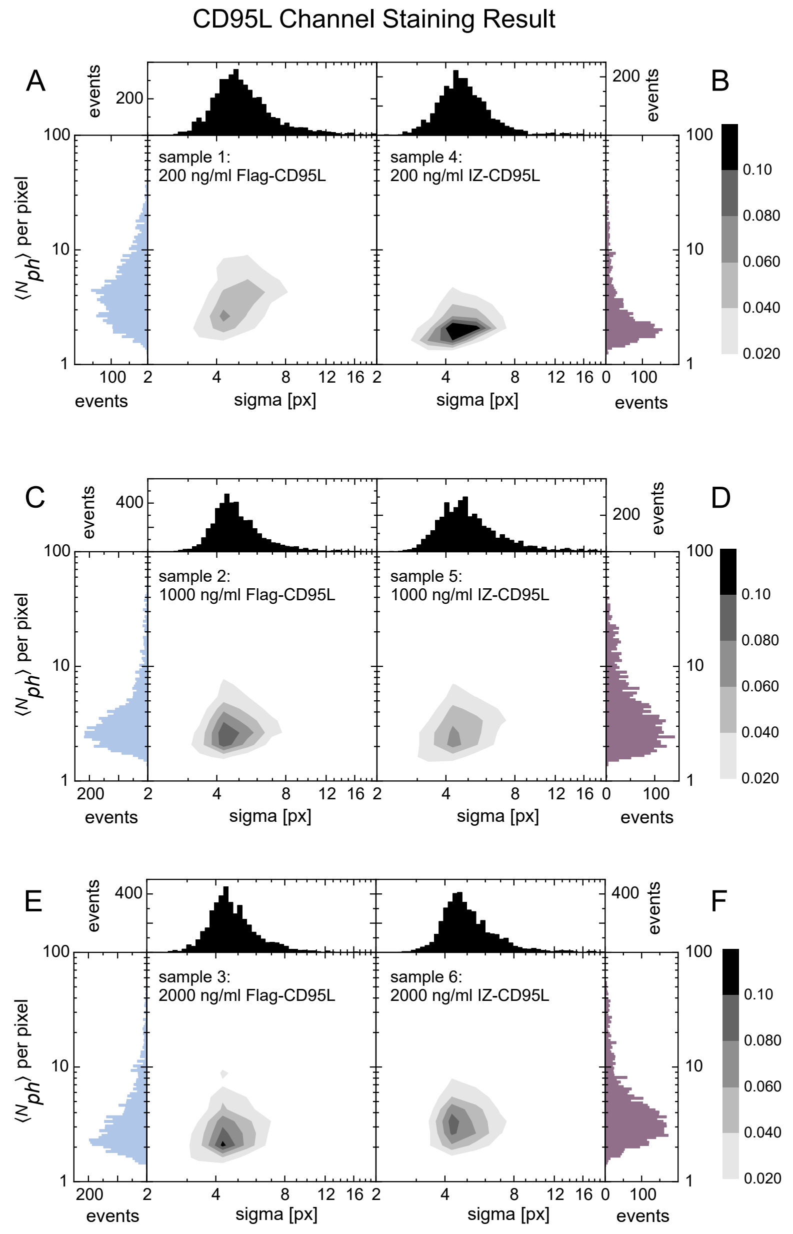


**Fig. S2.** CD95L results from quantitative spots analysis of two color stained STED images. Flag-CD95L or biotinylated IZ-CD95L at varying concentrations from 200 ng/ml, 1000 ng/ml, to 2000 ng/ml were incubated with Hela WT cells for 2 h at R.T. Sample preparation and data analysis was the same as in Fig.S1. Despite the increasing concentration, the ligand exhibits narrow distributions with small sigma $\sim$ 4 [px] and low brightness $N_{\mathrm{ph}}$ per pixel values $\sim$ 2-3. Only a slight non-significant increase in brightness change for IZ-CD95L is obtained. All distributions primarily correspond to the detection of single (oligomeric) ligands within spots. While no direct absolute value comparison of the CD95L channel with the CD95 channel is possible due to the different antibody and fluorophore, the distribution here suggests that only single oligomerization of ligand can be detected.

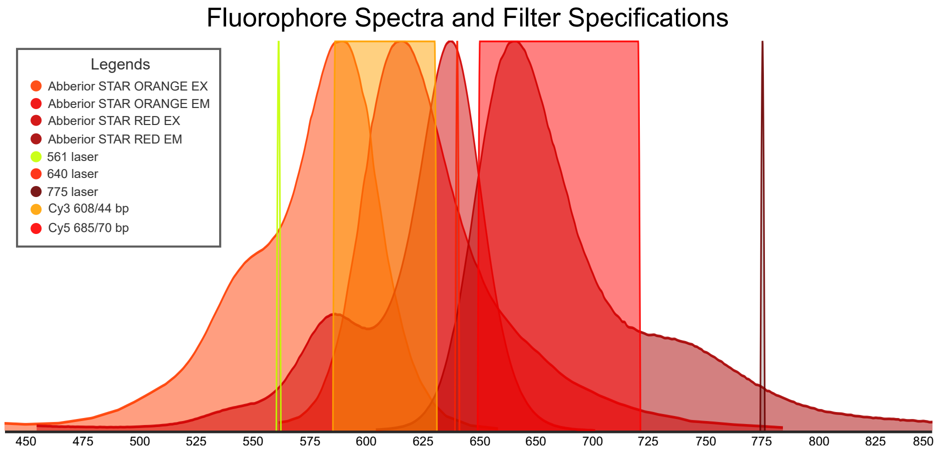


**Fig. S3.** Emission and excitation spectra of Abberior STAR dyes with laser lines and filters specifications. The figure was taken from FPbase::(https://www.fpbase.org/). In the Abberior STED microscope a single depletion laser at 775 nm is used for both excitation lasers at 561 nm and 640 nm. The Cy3 filter cube (Abberior ET608/44) and Cy5 filter cube (Abberior ET685/70M) were used for fluorescence detection of abberior STAR ORANGE (AbSTAROrange) and abberior STAR RED (AbSTARRed) dye, respectively.

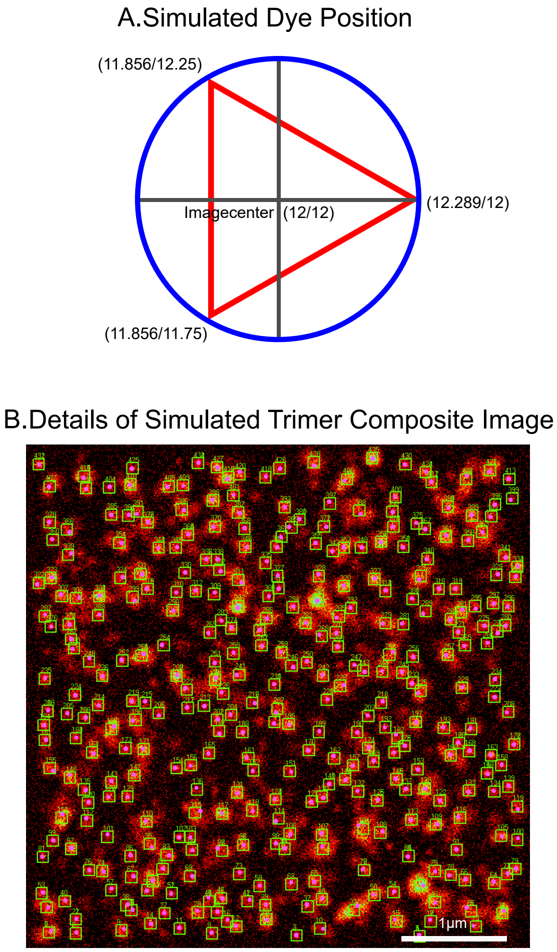


**Fig. S4.** Supporting Information for STED image simulation. A: Simulated dye positions of CD95 trimer. Three Gaussian distributions were placed at three defined positions in a ROI: (12.289, 12), (11.856, 12.25), (11.856, 11.75) with the image center located at (12, 12). CD95 dimers were positioned at identical intermolecular distances as in case of the trimer. CD95 monomers were positioned at the image center. B: Example of a composite image of 1000 simulated trimer ROIs randomly placed over the whole image. Scale bar: 1 µm.

**Table S1. Sample preparation and staining antibodies of Fig.5**

| Samples | Apoptosis Inducer | ligand concentration | Primary antibody | Secondary antibody |
|---|---|---|---|---|
| 1 | - | 200 ng/ml | $\alpha$-CD95 (Invitrogen, clone: JJ0942) | AbSTAROrange |
| 2 | Biotin-IZ-CD95L | 200 ng/ml | $\alpha$-CD95 (Invitrogen, clone: JJ0942) | AbSTAROrange |
| 3 | Flag-CD95L | 200 ng/ml | $\alpha$-CD95 (Invitrogen, clone: JJ0942) | AbSTAROrange |
| 4 | mAb-Flag-CD95L | 200 ng/ml | $\alpha$-CD95 (Invitrogen, clone: JJ0942) | AbSTAROrange |

**Table S2. Sample preparation and staining antibodies of Fig. S1 and S2**

| Samples | Apoptosis Inducer | ligand concentration | Primary antibody | Secondary antibody |
|---|---|---|---|---|
| 1 | Flag-CD95L | 200 ng/ml | $\alpha$-CD95L (Abcam), $\alpha$-CD95 (Biotium, B-R18) | AbSTAROrange, AbSTARRed |
| 2 | Flag-CD95L | 1000 ng/ml | $\alpha$-CD95L (Abcam), $\alpha$-CD95 (Biotium, B-R18) | AbSTAROrange, AbSTARRed |
| 3 | Flag-CD95L | 2000 ng/ml | $\alpha$-CD95L (Abcam), $\alpha$-CD95 (Biotium, B-R18) | AbSTAROrange, AbSTARRed |
| 4 | Biotin-IZ-CD95L | 200 ng/ml | $\alpha$-CD95L (Abcam), $\alpha$-CD95 (Biotium, B-R18) | AbSTAROrange, AbSTARRed |
| 5 | Biotin-IZ-CD95L | 1000 ng/ml | $\alpha$-CD95L (Abcam), $\alpha$-CD95 (Biotium, B-R18) | AbSTAROrange, AbSTARRed |
| 6 | Biotin-IZ-CD95L | 2000 ng/ml | $\alpha$-CD95L (Abcam), $\alpha$-CD95 (Biotium, B-R18) | AbSTAROrange, AbSTARRed |

**Table S3. Simulation parameters used in Fig.S4**

| Parameter name | Monomer | Dimer | Trimer |
|---|---|---|---|
| Amplitude of 2D Gaussians | $2.5 \pm 100\%$ [ph] | $2.5 \pm 100\%$ [Nph]<br>$2.5 \pm 100\%$ [Nph] | $2.5 \pm 100\%$ [Nph]<br>$2.5 \pm 100\%$ [Nph]<br>$2.5 \pm 100\%$ [Nph] |
| Poisson noise | yes | yes | yes |
| Gaussian position (X/Y) coordinates in $25 \times 25$ image | (12.000/12.000) | (12.000/12.250)<br>(12.000/11.750) | (11.856/12.250)<br>(11.856/11.750)<br>(12.289/12.000) |
| $\sigma_X = \sigma_Y$ of 2D Gaussians | $3.5 \pm 30\%$ [pixel] | $4.5 \pm 30\%$ [pixel]<br>$4.5 \pm 30\%$ [pixel] | $4.5 \pm 30\%$ [pixel]<br>$4.5 \pm 30\%$ [pixel]<br>$4.5 \pm 30\%$ [pixel] |
| Mean of Poisson background noise | 0.295 [Nph] | 0.295 [Nph] | 0.295 [Nph] |
| Composite image size | $500 \times 500$ [pixel] | $500 \times 500$ [pixel] | $500 \times 500$ [pixel] |
| Total number of generated spots | 1000 | 1000 | 1000 |
